# A conserved germline-specific Dsn1 alternative splice isoform supports oocyte and embryo development

**DOI:** 10.1101/2024.04.17.589883

**Authors:** Jimmy Ly, Cecilia S. Blengini, Sarah L. Cady, Karen Schindler, Iain M. Cheeseman

## Abstract

Alternative mRNA splicing can generate distinct protein isoforms to allow for the differential control of cell processes across cell types. However, alternative splice isoforms that differentially modulate distinct cell division programs have remained elusive. Here, we demonstrate that mammalian germ cells express an alternate mRNA splice isoform for the kinetochore component, DSN1, a subunit of the MIS12 complex that links the centromeres to spindle microtubules during chromosome segregation. This germline DSN1 isoform bypasses the requirement for Aurora kinase phosphorylation for its centromere localization due to the absence of a key regulatory region allowing DSN1 to display persistent centromere localization. Expression of the germline DSN1 isoform in somatic cells results in constitutive kinetochore localization, chromosome segregation errors, and growth defects, providing an explanation for its tight cell type-specific expression. Reciprocally, precisely eliminating expression of the germline DSN1 splice isoform in mouse models disrupts oocyte maturation and early embryonic divisions coupled with a reduction in fertility. Together, this work identifies a germline-specific splice isoform for a chromosome segregation component and implicates its role in mammalian fertility.

## Results and Discussion

### Identification of an evolutionarily-conserved germline-specific DSN1 splice isoform

The chromosome segregation and cell division programs associated with somatic mitosis and germline meiosis display dramatic differences [1–6]. The changes in chromosome segregation during meiosis require alterations to the established mitotic cell division machinery [5, 6]. However, it remains unclear what aspects of kinetochore function and its regulatory control differ between the mitotic and meiotic cell divisions to rewire these core processes. To understand the mechanisms that distinguish mitosis and meiosis, we sought to evaluate changes in the gene expression for the molecular machinery that directs chromosome segregation. Prior work has identified specific examples of germline-specific gene expression for cell division components, including Aurora Kinase C (AURKC) and Meikin, amongst others [5–8]. To test for additional examples of meiosis-specific behaviors, we compared mRNA-sequencing data from mitotically-arrested HeLa and human testes—a tissue rich in meiotically-dividing cells (Fig. 1A). On a gene level, we did not observe substantial differences in the expression levels for core kinetochore components between these cell types with the exception of these previously described examples (Fig. 1A and S1A). Thus, we next considered potential changes in the expression of mRNA isoforms for kinetochore components. To do this, we quantified the expression of individual mRNA isoforms in mitotically-arrested HeLa cells, whole human testes, and somatic human testis cells (Sertoli, Leydig, and Peritubular cells; Fig. 1B, 1C, and S1B). Of the mRNA isoforms that are enriched in whole testes, but not mitotic HeLa cells or somatic testes cells, we identified germline-specific transcriptional isoforms of the kinetochore proteins KNSTRN/SKAP [9] and MAD2L1BP/p31 Comet, each of which use an alternative upstream transcriptional start site (Fig. 1D). Finally, we observed a germline-specific alternate mRNA splice isoform for the kinetochore protein DSN1. This DSN1 isoform is the only testes-specific alternative splicing event observed for a kinetochore component identified in our analysis.

**Figure 1.**
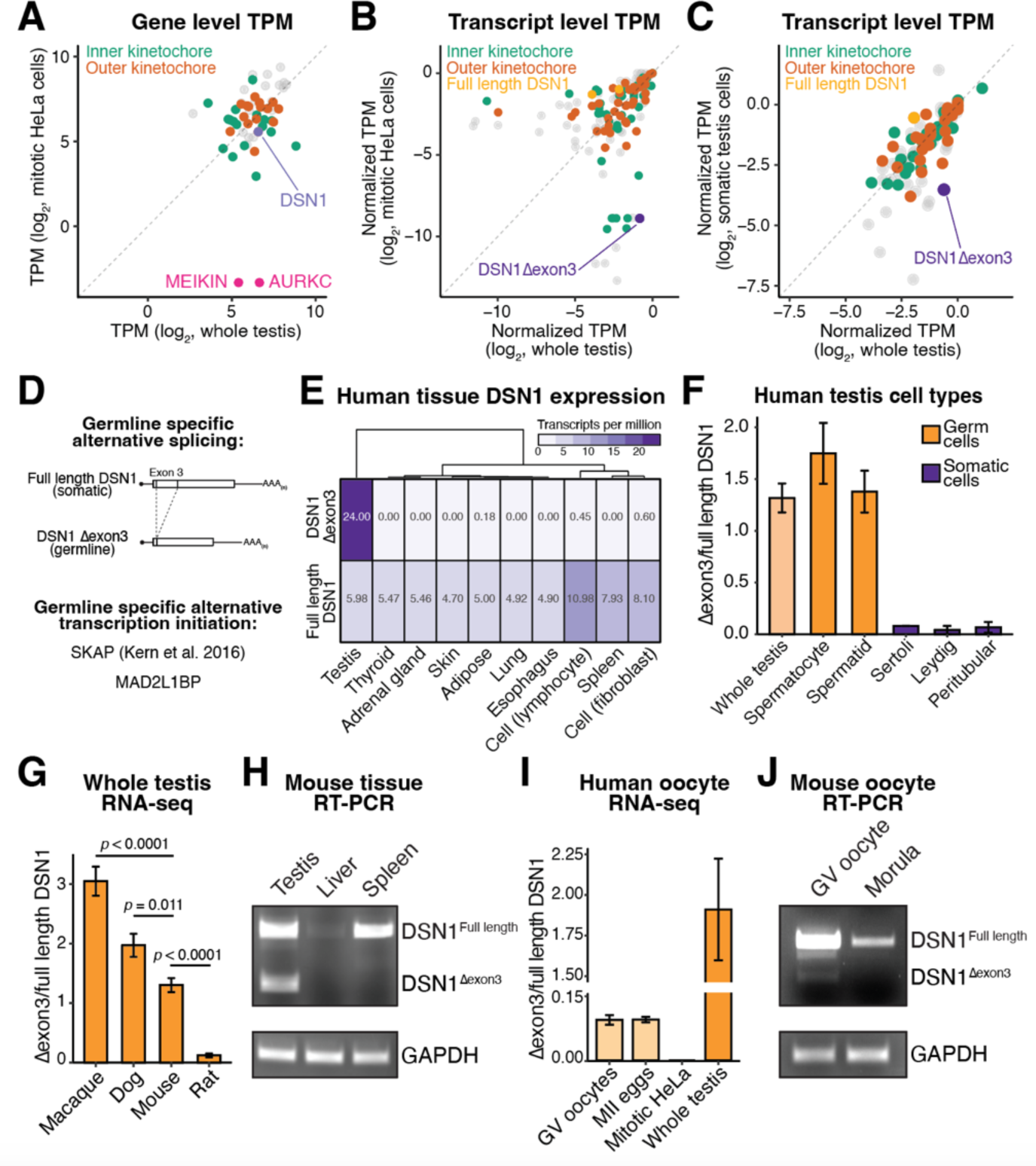
Identification of an evolutionarily-conserved germline specific DSN1 splice isoform. (**A**) Gene level mRNA expression for core kinetochore components in mitotically arrested HeLa cells and whole human testis. (**B**) Transcript level mRNA expression for core kinetochore components in mitotically arrested HeLa cells and whole human testis. (**C**) Transcript level mRNA expression for core kinetochore components in somatic testis cells (Leydig, Sertoli, and Peritubular cells) and whole human testis. (**D**) Alternative mRNA isoforms that are at least enriched 2-fold in whole testis compared to both mitotic HeLa and somatic testis. (**E**) DSN1 mRNA isoform expression in humans from the GTEx consortium. (**F**) DSN1 Δexon3/full length ratio in specific cells from human testis mRNA sequencing. Error bars indicate standard error of the mean and N = 3 biological replicates. (**G**) DSN1 Δexon3/full length ratio across organisms from mRNA sequencing data. Error bars indicate standard error of mean and N = 8 biological replicates, except for rat, which had 7. Student’s T-test was used to perform statistical test. (**H**) RT-PCR from mouse testis, liver, and spleen. DSN1 Δexon3 is detected specifically in mouse testis. (**I**) DSN1 Δexon3/full length ratio in human oocytes, metaphase II eggs, mitotic HeLa, and whole testis in mRNA sequencing. DSN1 Δexon3 is detected in female germline. Error bars indicate standard error of the mean. N = 10 biological replicates for GV oocytes and MII eggs (5 young and 5 advanced maternal age female donors). N = 3 biological replicates for mitotic HeLa cells and whole testis. (**J**) RT-PCR from mouse GV oocytes and morula. DSN1 Δexon3 is detected specifically in mouse GV oocytes and is undetected in morula.

DSN1 is a subunit of the 4-protein Mis12 complex, which interacts with the inner kinetochore protein, CENP-C, and recruits the NDC80 complex to centromeres to direct interactions with spindle microtubules [10, 11]. This DSN1 splice isoform removes exon 3 of the mRNA, resulting in a 106 amino acid internal deletion (referred to here as DSN1^Δexon3^). DSN1^Δexon3^ was specific to the human testes and was not detected appreciably in other human tissues based on analysis of hundreds of patient samples from the GTEx consortium (Fig. 1E, [12]). Analysis of RNA-seq data from spermatocytes, mature spermatids, Leydig, peritubular, and Sertoli cells revealed that, within the testis, DSN1^Δexon3^ is primarily expressed in meiotically-dividing cells (Fig. 1F; [13, 14]). The germline-specific expression of the DSN1^Δexon3^ splice isoform was conserved across multiple mammals including macaque, dog, mouse, and rats (Fig. 1G; [15]), but with rodents expressing a decreased amount of the germline DSN1^Δexon3^ isoform relative to the canonical isoform in testes. We note that this difference in DSN1 isoform ratio between organisms is unlikely to be due to differences in the relative ratio of meiotic/mitotic cells in the testis between organisms as measured based on relative *REC8*, *KIF2B*, *AURKC*, and *MEIKIN* mRNA levels (Fig. S1D). In addition to the analysis of public datasets, we validated the testis-specific expression of DSN1^Δexon3^ in mouse testis using RT-PCR (Fig. 1H and S1C). We also detected DSN1^Δexon3^ in mouse oocytes, indicating that this splicing event is germline-specific rather than male-specific (Fig. 1I and 1J; [16]). However, we note that the relative abundance of germline DSN1^Δexon3^ compared to the full length DSN1 isoform is higher in the male compared to the female germline. Together, these data suggest that germ cells express an evolutionarily-conserved mRNA splice isoform of DSN1 lacking exon 3.

### The germline-specific DSN1 splice isoform bypasses the requirement for aurora kinases for centromere localization

The germline-specific splice isoform of DSN1 identified above results from an exon skipping event that removes exon 3. Exon 3 encodes amino acids 12-118, which includes a critical regulatory region of DSN1 (amino acids 100-109) that is subject to phosphorylation by the kinase Aurora B at Serine 100 and 109 [17–19]; Fig. 2A and 2B). This DSN1 phosphorylation acts to regulate the interaction between the MIS12 complex and CENP-C in a cell cycle-dependent manner [17–21]. To test the consequences of eliminating the amino acids encoded by exon 3, we first tested the ability of DSN1^Δexon3^ to localize to centromeres during metaphase in somatic cells. Ectopic expression of a GFP fusion with the germline DSN1^Δexon3^ isoform in HeLa cells displayed kinetochore localization during metaphase similar to somatic full length DSN1 (Fig 2C). Moreover, based on quantitative IP-mass spectrometry of GFP tagged germline DSN1^Δexon3^, DSN1^Δexon3^ was able to interact with core kinetochore components when expressed in somatic cells (Fig. 2D; Table S1). These results indicate that the removal of Dsn1 exon 3 (residues 12-118) does not disrupt the ability of Dsn1 to act as an integral subunit of the Mis12 complex at kinetochores.

**Figure 2.**
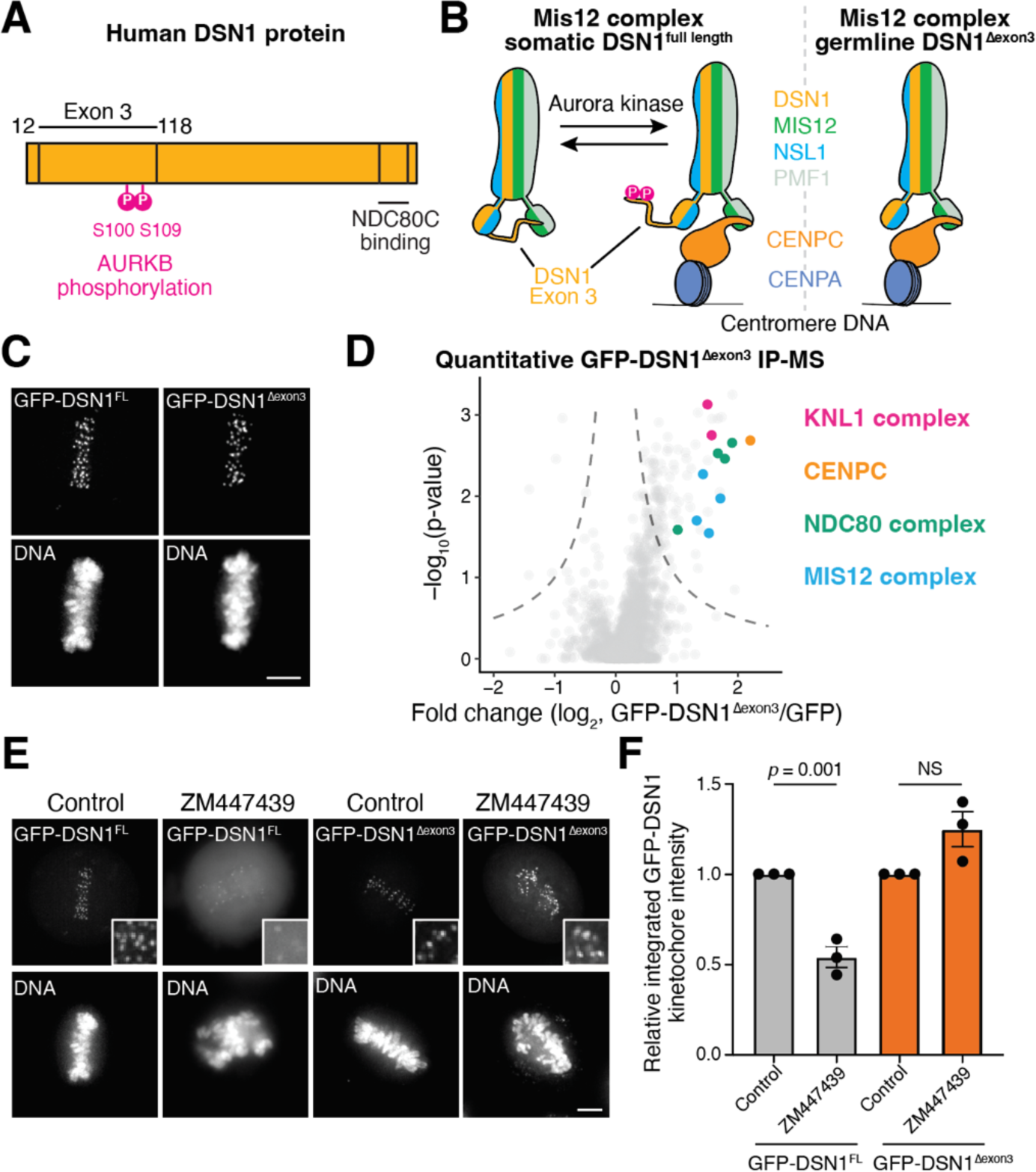
Germline specific DSN1 isoform bypasses the requirement for AURKC for centromere localization. (**A**) Schematic representation of full length DSN1. Exon 3 of DSN1 encodes for a regulatory region of DSN1 (amino acids 12-118) harboring the phosphorylation sites (S100 and S109) for Aurora kinases. (**B**) Schematic representation of the role of DSN1 exon 3 in regulating centromere localization. DSN1 is a member of a 4 subunit MIS12 protein complex containing DSN1, MIS12, NSL1, and PMF1. Full length DSN1 cycles between an open and closed conformation. DSN1 exon 3 autoinhibits MIS12 complex centromere localization by interacting with the surface that interacts with CENP-C. Phosphorylation by Aurora kinases relieves this autoinhibition allowing for the MIS12 complex to interact with CENP-C. DSN1 lacking exon 3 deletes the autoinhibitory region thus allowing constitutive centromere localization. (**C**) Live cell imaging showing the localization of ectopically expressed GFP-DSN1 full length and GFP-DSN1^Δexon3^. Both constructs are able to localize to centromeres during interphase. (**D**) Volcano plot from quantitative mass spectrometry of GFP-DSN1^Δexon3^ compared to GFP immunoprecipitations. GFP-DSN1^Δexon3^ is able to interact with the core kinetochore complexes, MIS12, NDC80, KNL1, and CENP-C. (**E**) Live cell imaging of GFP-DSN1 full length and GFP-DSN1^Δexon3^ in the absence or presence of the Aurora kinase inhibitor, ZM447439. (**F**) Quantification of GFP-DSN1 kinetochore intensity from (E). Error bars indicate standard error of the mean and students T test was used for statistical test. N = 3 biological replicates, which contained >10 cells per replicate, and >20 kinetochores per cell.

When dephosphorylated, the N-terminal region of DSN1 acts in an autoinhibitory manner to prevent the MIS12 complex from binding to CENP-C thus blocking its centromere localization ([20, 21]; Fig. 2B). Phosphorylation of DSN1 by Aurora B inhibits the ability of this N-terminal region to act in an autoinhibitory manner thereby promoting the interaction of the Mis12 complex with CENP-C and its recruitment to centromeres. Importantly, although the localization of somatic DSN1 was sensitive to Aurora B inhibition, the localization of ectopically-expressed GFP-tagged germline DSN1^Δexon3^ to centromeres was unchanged even when Aurora B was inhibited by treatment with ZM447439 (Fig. 2E, 2F; Fig. S2A). Together, our results suggest that germline DSN1^Δexon3^ is able to interact and localize similarly to somatic DSN1, but bypasses the requirement of phosphorylation for its recruitment to centromeres.

### The germline-specific DSN1 splice isoform localizes to kinetochores throughout the cell cycle

Under normal conditions, the outer kinetochore assembles immediately prior to mitotic entry and disassembles at the end of mitosis [22]. Since the phospho-regulation of DSN1 acts to regulate Mis12 complex centromere recruitment, we next tested the cell cycle-dependent localization of DSN1^Δexon3^. Full-length DSN1 delocalized from kinetochores upon anaphase onset (Fig. 3A; [18, 19, 23]). In contrast, we observed centromere localization for GFP-DSN1^Δexon3^ in anaphase and early G1 cells (Fig. 3A). This prolonged localization for Dsn1^Δexon3^ persisted throughout the cell cycle as every interphase cell observed displayed centromere-localized DSN1 (Fig. 3B). Similarly, when expressed ectopically in HeLa cells, GFP-tagged mouse DSN1^Δexon3^, which displays 70% sequence identity to the human protein, also persisted at kinetochores throughout the cell cycle (Fig. S2A-C), suggesting a similar regulatory control. Consistent with the constitutive interphase centromere localization, immunoprecipitations and quantitative mass spectrometry revealed that germline DSN1^Δexon3^ is more strongly associated with its direct centromere binding partner CENP-C during interphase compared to somatic full-length DSN1 (Fig. 3C; Table S1). These data suggest that germline DSN1^Δexon3^ displays distinct regulatory control enabling it to persist at kinetochores throughout interphase (Fig. 2B).

**Figure 3.**
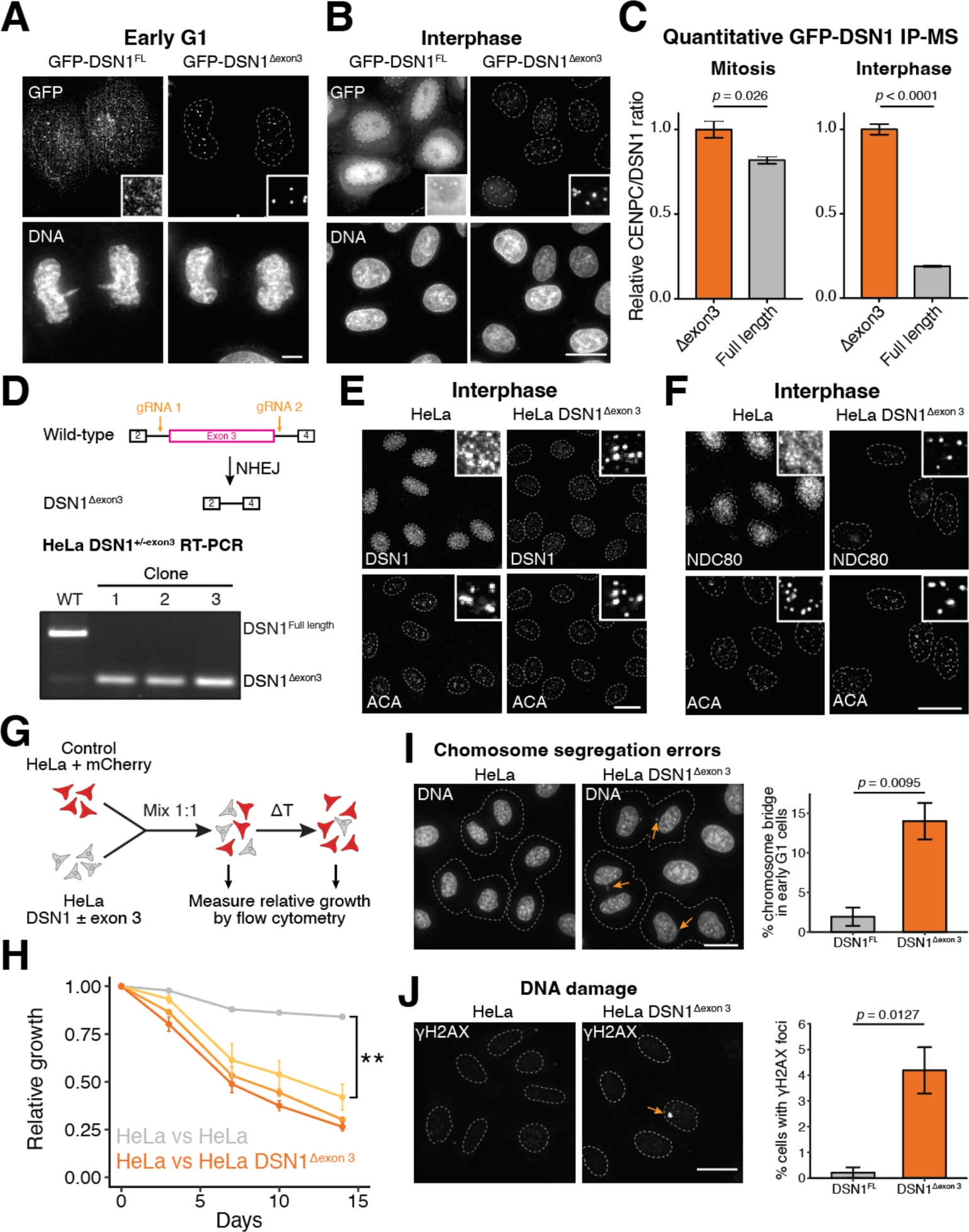
The germline-specific DSN1 splice isoform localizes to centromeres throughout the cell cycle and is deleterious to somatic cells. (**A**) Live cell imaging of GFP-DSN1 full length and GFP-DSN1^Δexon3^ in late anaphase. GFP-DSN1^Δexon3^ persists at centromeres while GFP-DSN1 full length is evicted. Scale bar represents 5 µM. (**B**) Live cell imaging of GFP-DSN1 full length and GFP-DSN1^Δexon3^ during interphase. GFP-DSN1^Δexon3^ is still able to localize to centromeres. Scale bar represents 20 µM. (**C**) Barplot showing the relative CENP-C interaction from GFP-DSN1 quantitative IP-MS experiments. Both GFP-DSN1 full length and GFP-DSN1^Δexon3^ strongly interact with CENP-C during mitosis but only GFP-DSN1^Δexon3^ is able to strongly associate with CENP-C during interphase. Error bars indicate standard error of the mean and students T test was used for statistical test. (**D**) Top, schematic representation of gene editing strategy to endogenously remove exon 3 from DSN1. Bottom, RT-PCR analysis using primers flanking exon 3 of wild type HeLa and 3 different clones of HeLa^Δexon3^. (**E**) Immunofluorescence images of DSN1 during interphase in control HeLa and HeLa^Δexon3^ cells. Scale bar represents 20 µM. (**F**) Immunofluorescence images of NDC80 during interphase in control HeLa and HeLa^Δexon3^ cells. Scale bar represents 20 µM. Scale bar represents 20 µM. (**G**) Schematic outline of competitive growth assays. Control mCherry HeLa cells are mixed with control or Δexon3 HeLa cells and their relative growth is monitored by flow cytometry. (**H**) Competitive growth assay suggests that HeLa cells lacking Δexon3 have growth defects. 3 clonal HeLa^Δexon3^ (different clones in different shades of orange) were tested. N = 3 biological replicates and error bars indicate standard error of the mean. Student’s T test was used for statistical test, ** p < 0.005. (**I**) Left, Immunofluorescence images of DNA indicates increased chromosome segregation errors in HeLa^Δexon3^ cells. Scale bar represents 20 µM. Right, quantification of the number of cells with chromosome segregation errors. N = 3 biological replicates, error bars indicate standard error of the mean, and students T test was used for statistical test. (**J**) Left, Immunofluorescence images of γH2AX indicates increased DNA damage in HeLa^Δexon3^ cells. Scale bar represents 20 µM. Right, quantification of the number of cells with bright γH2AX foci. N = 3 biological replicates, error bars indicate standard error of the mean, and students T test was used for statistical test.

### Somatic cells do not tolerate constitutive germline-specific DSN1 expression

Given the critical role of this Dsn1 regulatory region in controlling kinetochore assembly, we hypothesized that there may be functional consequences of expressing germline DSN1^Δexon3^ in somatic cells. To test this, we eliminated exon 3 from the endogenous DSN1 locus in somatic cells using Cas9-based gene editing (Fig. 3D). Removal of this exon resulted in cell lines that only express the germline DSN1^Δexon3^ isoform (Fig. 3D). Similar to ectopically-expressed GFP-DSN1 fusion proteins (Fig. 2E, 3A, and 3B), we found that endogenous germline DSN1^Δexon3^ was insensitive to Aurora B inhibition and localized to centromeres throughout the cell cycle (Fig. 3E, 3F; Fig. S2D, S3A-C).

We next measured the growth of HeLa DSN1^Δexon3^ cells using competitive growth assays (Fig. 3G). Although DSN1^Δexon3^ cells were viable, we observed a decrease in growth of DSN1^Δexon3^ cells compared to control HeLa cells (Fig. 3H). In addition, DSN1^Δexon3^ cells displayed an increased frequency of chromosome bridges during anaphase (Fig. 3I). Consistent with increased frequency of chromosome bridges, DSN1^Δexon3^ cells had an increased prevalence of interphase γH2AX foci, a marker of DNA damage (Fig. 3J). These data suggest that unresolved interphase DNA damage may be one mechanism to explain the increased incidence of chromosome segregation errors. Together, our results suggest that proper outer kinetochore disassembly is required for robust somatic cell growth and shed light on why the expression of the germline DSN1^Δexon3^ splice isoform is tightly restricted to the germline.

### The germline DSN1^Δexon3^ splice isoform is required for robust meiosis, early embryonic development, and fertility

DSN1^Δexon3^ is expressed specifically in mouse testis and oocytes (Fig. 1). Following meiosis I in mammals, cells immediately enter meiosis II without an intervening growth or DNA replication phase. The short timing between these meiotic divisions and the requirement for assembled kinetochores for both of these stages suggests that meiotic cells may require persistent stable outer kinetochore formation. Since the germline DSN1^Δexon3^ isoform promotes stable kinetochore assembly even after chromosome segregation, we hypothesized that this splice isoform may contribute to robust meiotic divisions (Fig. S3D). To test this, we generated a mouse model in which we eliminated the production the germline-specific DSN1 splice isoform (Fig. 4A and 4B). To precisely block the production of germline DSN1^Δexon3^, we deleted the intron between exon 2 and exon 3 to prevent the removal of exon 3, thereby enforcing the exclusive production of full length DSN1 in the germline (DSN1 Δintron2, Fig. 4A-C). To generate this mouse model, we used CRISPR/Cas9 to eliminate DSN1 intron 2 in mouse embryonic stem cells, injected the engineered embryonic stem cells into blastocysts, and pseudo-implanted the chimeric blastocysts into a donor female mouse. Animals were backcrossed at least 10 times to C57BL/6J animals to ensure a clean genetic background. DSN1 Δintron2/Δintron2 animals were viable with no apparent health issues (data not shown), consistent with DSN1^Δexon3^ only being expressed in the germline.

**Figure 4.**
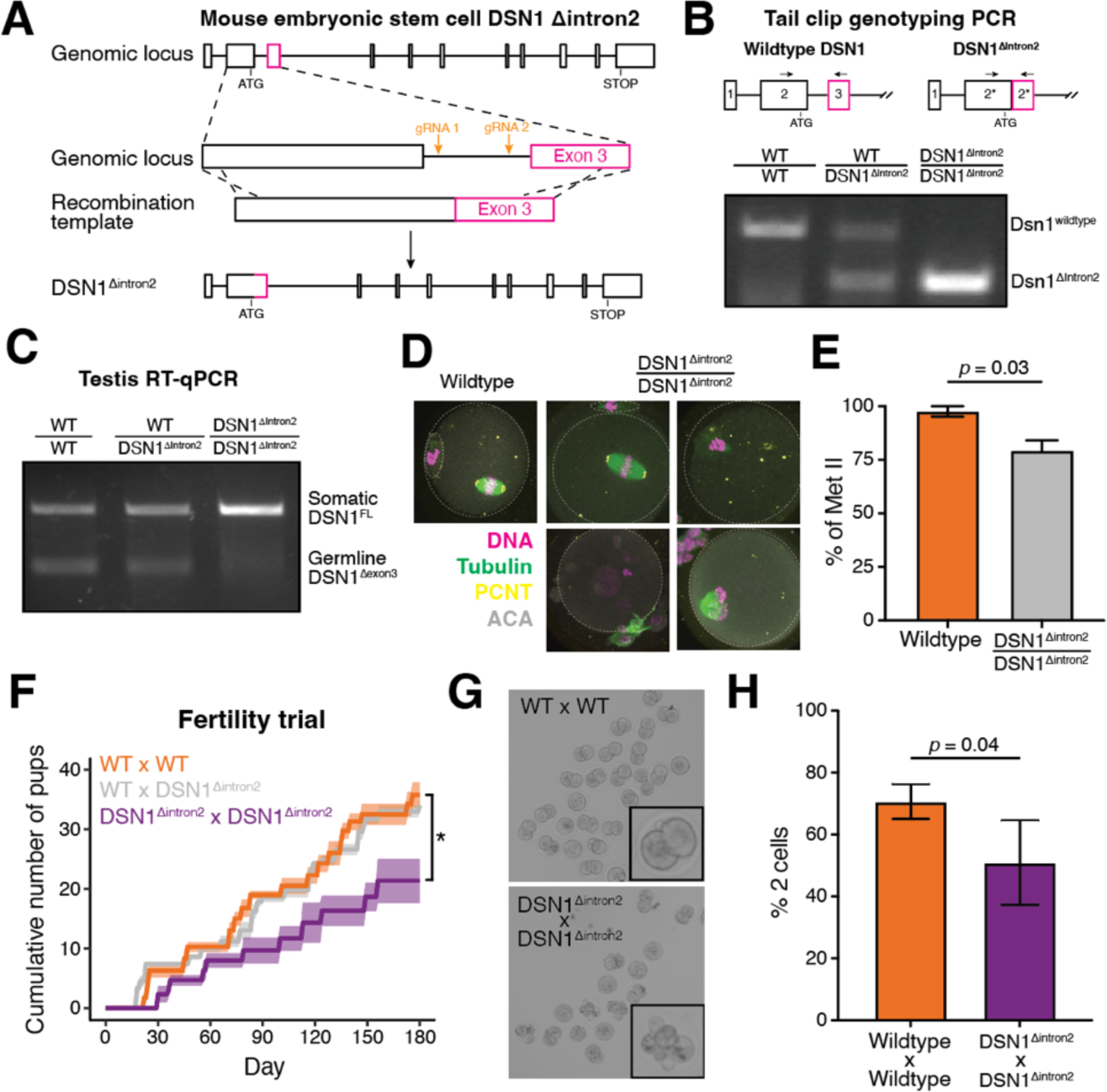
Germline DSN1 splice isoform is required for robust oocyte and embryo development and contributes to fertility. (**A**) Schematic outline of DSN1 Δintron2 mouse model generation. DSN1 locus in mouse embryonic stem cells was editing using CRISPR/Cas9 to remove intron 2, therefore inhibiting removal of exon 3. DSN1 Δintron2 mouse embryonic stem cells were used to establish mouse model. (**B**) Genotyping agarose gel of PCRs from tail clips from the indicated mouse genotypes. (**C**) RT-PCR analysis using primers flanking mouse DSN1 exon 3 of mouse testis from indicated genotypes. (**D**) Representative images of metaphase II eggs from indicates genotypes. (**E**) Quantification of normal metaphase II from indicated mouse genotypes in (D). N = 3 biological replicates. Error bars indicate standard error of the mean and student’s T-test was used for statistical test. (**F**) 6-month fertility assay from indicated genotype. Line indicates average number of pups and shade represents standard error of mean between biological replicates. N = 4 for wildtype female x wildtype male, N = 5 for DSN1 Δintron2/Δintron2 female x wildtype male, N = 3 for DSN1 Δintron2/Δintron2 female x DSN1 Δintron2/Δintron2 male. Student’s T test was used for statistical test, * p < 0.05. (**G**) Representative images of 2 cell embryos from indicated mating. (**H**) Quantification of the number of 2-cell embryos in (F). N = 4 for WT x WT and N = 3 for DSN1 Δintron2/Δintron2 x DSN1 Δintron2/Δintron2 mating. Error bars indicate standard error of the mean and student’s T-test was used for statistical test.

To test if germline DSN1 splicing affects oocyte quality, we first isolated metaphase II eggs from wild type (C57BL/6J) and DSN1 Δintron2/Δintron2 homozygous females and analyzed chromosome alignment. We observed an increased number of chromosome alignment errors in DSN1 Δintron2/Δintron2 female metaphase II eggs (Fig 4D and 4E). We next considered the possibility that DSN1 Δintron2/Δintron2 animals may be sub-fertile. Over the course of a 6-month fertility assay, we did not observe a strong change in the cumulative litter size when mating DSN1 Δintron2/Δintron2 females to wild-type males (Fig. 4F). In contrast, we observed a modest reduction in the overall number of newborn animals in matings between DSN1 Δintron2/Δintron2 females and DSN1 Δintron2/Δintron2 males. Together, our results suggest that the germline-specific DSN1^Δexon3^ alternative splice isoform is important for robust fertility and oocyte development.

We next asked why the fertility defects were observed when we crossed DSN1 Δintron2/Δintron2 females to DSN1 Δintron2/Δintron2 males, but not to wild-type males. As any paternal contribution must occur post-fertilization, we therefore assessed defects in early embryonic development in DSN1 Δintron2/Δintron2 animals. To test embryonic development, we mated DSN1 Δintron2/Δintron2 female mice to DSN1 Δintron2/Δintron2 males and isolated 1-cell embryos and matured them *in vitro* to measure embryonic development. We found that DSN1 Δintron2/Δintron2 females crossed to DSN1 Δintron2/Δintron2 males gave rise to an increased number of defective embryos (Fig. 4G and 4H). In particular, an increased proportion of these DSN1 Δintron2/Δintron2 embryos were stuck in the 1-cell embryo stage or died, whereas control embryos developed normally to the 2-cell stage (Fig. 4G and 4H). However, for DSN1 Δintron2/Δintron2 embryos that survived to the 2-cell stage, most of them developed normally to the morula stage (Fig. S4A and S4B). This suggests paternally contributed DSN1^Δexon3^ is required for robust early embryonic development.

This apparent role for germline DSN1^Δexon3^ in early embryos was unanticipated given our initial gene expression analysis suggesting meiosis-specific DSN1^Δexon3^ expression. However, DSN1^Δexon3^ may not be cleared in the early embryos, or it may also be transcribed during early embryonic divisions. To test this, we measured the expression of germline DSN1^Δexon3^ during the course of early embryonic development by analyzing previously published embryo RNA-seq data from human [24] and mouse [25]. We were able to detect DSN1^Δexon3^ in early embryos prior to zygotic transcription (Fig. S4E and S4F). Moreover, we observed strong DSN1^Δexon3^ expression near the onset of zygotic transcription (8-cell stage in humans and 1 to 2-cell stage in mice) that is cleared in the following cell divisions (Fig. S4E and S4F). The expression of DSN1^Δexon3^ in early embryos along with its zygotic expression are consistent with a role for DSN1^Δexon3^ in early mammalian embryos in addition to its role during oocyte meiosis. Together, our results suggest that the germline-specific DSN1^Δexon3^ is required for both oocyte meiosis and early embryo divisions and that this splice isoform is important for robust fertility.

### A conserved germline-specific DSN1 alternative splice isoform promotes stable kinetochore assembly and is required for early development

Here, we identify the first reported mammalian germline-specific splice isoform for a kinetochore component. We demonstrate that this germline-specific DSN1 isoform bypasses the requirement for Aurora B phosphorylation for centromere localization, resulting in persistent outer kinetochore localization even following chromosome segregation when expressed in somatic cells. This germline Dsn1 splice isoform is required for robust meiotic division in mouse models, particularly in metaphase II. These germline defects may reflect an altered regulatory control of kinetochore assembly, error correction, or meiotic arrest behavior. Moreover, we implicate this germline DSN1^Δexon3^ splice isoform in early embryonic development and fertility. We hypothesize that DSN1^Δexon3^ is present in early embryos and is transcribed in the early rounds of zygotic transcription to allow for transient DSN1^Δexon3^ expression, which contributes to early embryo development. We speculate that differences in early and late embryo cell divisions may place an enhanced and unanticipated requirement for the DSN1^Δexon3^ splice isoform during the early embryonic divisions. Although the early embryonic cell cycle requires DNA replication and intervening gap-phases, the early embryo cell cycle is distinct from a late embryo cell cycle. For example, early and late embryo cell cycles differ in the length of gap-phases, timing and length of mitosis, amount of aneuploidy, and the presence of DNA damage [26–28]. These differences in the cell cycle may place a requirement on prolonged outer kinetochore assembly or the altered regulation of error correction during early embryo development through the activity of DSN1^Δexon3^. Alternatively, DSN1^Δexon3^ may play other unanticipated roles during the early embryonic divisions that are required for development.

Although the reduction in oocyte and embryo quality along with the observed fertility defects in DSN1 Δintron2/Δintron2 animals are reproducible and statistically significant, this represents an intermediate effect and the mutant mice do not display infertility. We speculate that germline DSN1^Δexon3^ may become particularly important in different genetic backgrounds, physiological conditions, or organisms. In particular, some mammals, such as primates or dogs, display a much higher ratio of germline vs. somatic DSN1 in the germline (Fig. 1G), which may suggest an enhanced requirement in these organisms. Moreover, compensation through other outer kinetochore recruitment pathways may contribute to the modest effects—such as through compensation from the CENP-T [29] or AURKB pathways [30, 31]. In contrast to the need for the germline DSN1^Δexon3^ for proper fertility, we demonstrate that forced expression of germline DSN1 in somatic cells results in decreased cell fitness. Since the kinetochore composition appears normal in mitosis (Fig. S3E), we speculate that the observed DNA damage in these cells arise from problems with centromere DNA replication in the presence of constitutively assembled outer kinetochore [22].

## Acknowledgments

We thank the members of the Cheeseman lab, in particular Alexandra Nguyen for helpful discussions; the Whitehead Genetically Engineered Models Center for mouse model generation and the Whitehead Quantitative Proteomics Core for mass spectrometry on the Orbitrap Eclipse.

## Author contributions

Conceptualization: IMC, JL, CSB, and KS

Methodology: JL, CSB, and SLC

Investigation: CSB for mouse oocyte and embryo phenotyping, SLC for the mouse fertility trial and competition assay, JL all other experiments

Formal analysis: CSB for mouse oocyte and embryo phenotyping, SLC for the mouse fertility trial and competition assay, JL all other experiments

Writing: JL and IMC wrote the manuscript with input from KS

Supervision: IMC and KS

Funding acquisition: IMC, KS, and JL

## Experimental Procedures

### Genomes and RNA-seq analysis

Human and mouse transcriptome were downloaded from the GENCODE website (release 25, GRCh38.p7 for humans and release 25, GRCm38.p6 for mice). Crab-eating macaque, dog, and rat transcriptome were downloaded from the Ensembl website (Ensembl release 102 Macaca_fascicularis_5.0; Ensembl release 111 Ros_Cfam_1.0, and Ensembl release 111 Rat-SHRSP/BbbUtx(Rattus norvegicus)_1.0). A custom transcriptome was used for the mouse and rat, where DSN1^Δexon3^ transcript had to be added to the downloaded transcriptome because this DSN1^Δexon3^ transcript was not in the annotations.

The transcriptome was indexed using Kallisto [32] with default settings. The RNA-sequencing data was reanalyzed from [13, 14] (Human testis samples, GSE74896); [33] (Mitotic HeLa cell samples, GSE230189); [15] (Monkey, Dog, Mouse, and Rat samples, GSE125483); [16] (Human GV oocytes and MII eggs, GSE95477); and [24] (Human and mouse embryo time course samples, GSE71434). For all sequencing analysis, transcript level quantification was performed using Kallisto with default settings. For gene level analysis, the TPM for each mRNA isoform was summed for the sample.

For reanalysis of GTEx data, the top 3 most abundant DSN1 transcripts were summed to represent full length DSN1 (ENST00000373750, ENST00000480153, ENST00000448110) and compared to the single DSN1^Δexon3^ transcript (ENST00000373734).

### Tissue culture

HeLa cells were cultured in DMEM with 10% heat-inactivated fetal bovine serum, 2 mM L-glutamine and 100 U/mL penicillin-streptomycin at 37°C with 5% CO2. V6.5 mouse embryonic stem cells were cultured on irradiated MEFs in DMEM supplemented with 10% FBS, 20 µg/mL recombinant mouse leukemia inhibitory factor (mLIF), 0.1 mM beta-mercaptoethanol, 100 U/mL penicillin-streptomycin, 2 mM L-glutamine, and 1% nonessential amino acids. Embryonic stem cells were cultured on MEF feeder gelatin-coated plates.

### RNA isolation

Mouse tissues were flash frozen then stored at -80°C. Frozen tissue was homogenized using a mortar-pestle on dry ice to maintain the dissociated cells frozen. A small aliquot of frozen crushed up tissues was resuspended in 400uL TRI Reagent (Invitrogen, AM9738). Approximate 100 oocytes or embryos from multiple animals was resuspended 400uL TRI Reagent. HeLa cells were washed once with PBS, resuspended 400 µL of TRI Reagent, and stored at -80°C. 120 µL of chloroform was added to the TRI reagent, the mixture was vortexed then phase separated by centrifugation at 21000x g, 4°C for 15 minutes. The aqueous layer containing the nucleic acids was collected mixed with an equal volume of chloroform and centrifuged 21000x g, 4°C for 1 minute. The RNA was precipitated with 300mM NaCl and 30µg GlycoBlue (AM9516) by the addition of equal volumes of isopropanol and incubation at -20°C overnight. Precipitated RNA was collected by centrifugation at 21000x g, 4°C for 15 minutes. The supernatant was removed and the RNA pellet was washed with 70% ethanol then dissolved in RNase-free water. The RNA concentration was quantified by nanodrop.

### Reverse transcription and PCR

The Maxima First Strand cDNA Synthesis Kit for RT-qPCR (ThermoFisher, K1671) was used for DNase treatment and reverse transcription according to manufacturer’s protocol. Total cDNA was then used for PCR with the Q5 High-Fidelity 2X Master Mix according the manufacturer’s instructions.

### Molecular biology and human cell line generation

Human DSN1 (ENST00000373750), Human DSN1^Δexon3^ (ENST00000373734), Mouse DSN1 (ENSMUST00000103129), and Mouse DSN1^Δexon3^ (ENSMUST00000103129 but without exon 3, this isoform is not annotated in the Ensembl annotations) was cloned downstream of pBABEblast-GFP [34]. GFP-SPC24 was used from [35].

Retroviral plasmids (pBABEblast-GFP fusion protein) were cotransfected into HEK293-GP cells along with the VSV-G plasmid for the production of retrovirus. The resulting retroviral was incubated with HeLa or DSN1^Δexon3^ cells for 48 hours with 10 μg/ml polybrene. The media was replace then grown for 48 hours. Infected cells were sorted for GFP by flow cytometry using the same gate for all GFP-tagged constructs to generate a polyclonal population of cells expressing similar amounts of GFP-tagged proteins.

To generate DSN1^Δexon3^ cells, HeLa cells were co-transfected with pX330-sgRNA plasmids and mCherry-expressing plasmid using Lipofectamine 2000. The pX330-sgRNA plasmids consisted of two pairs of guide RNAs, each pair flanking exon 3. Cleavage guided by the pair of guide RNAs followed by non-homologous end joining results in the removal of exon 3. Following transfection, mCherry expressing cells were singly sorted into 96-well plates and grown up. Individual clones were genotyped using primers flanking exon 3 to ensure homozygous removal of DSN1 exon 3. Three independent clones were used in this manuscript, 2 using the first pair of sgRNA and 1 using the second pair of sgRNA. The first pair of guide RNA sequence: 5’-TCTTTATGTTTCCACGATCT-3’ and 5’-CGGTATCTACATCTTGCGAT-3’. The second pair of guide RNA sequence: 5’-TTTATGTTTCCACGATCTAG-3’ and 5’-TCTACATCTTGCGATTGGTT-3’.

### Immunoprecipitations

Polyclonal GFP antibodies was coupled to Protein A beads as described in [34]. For GFP-DSN1 IP experiments, 5 15cm plates of interphase cells were dislodged using PBS+ 5 mM EDTA resuspended in DMEM and spun down at 500 x g for 5 minutes. For mitotic cell enrichment, 8 15cm plates were treated with 8 µM STLC 16 hours, mitotic cells were shaken off and spun down at 500 x g for 5 minutes. Cell pellet was washed once in ice cold PBS then lysis buffer (50 mM HEPES pH7.4, 1 mM EGTA, 1 mM MgCl2, 300 mM KCl, 10% glycerol). The cell pellet was then resuspended in an equal volume of lysis buffer, flash frozen in liquid nitrogen, and stored at -80°C. Cells were thawed in a 37°C water bath with an equal volume of 1.5X lysis buffer supplemented with 0.075% Nonidet P-40, 1 mM phenylmethylsulfonyl fluoride, and cOmplete EDTA-free protease inhibitor cocktail (Roche, 11873580001). Cells were lysed by sonication and cleared by centrifugation at 21000x g for 30 minutes at 4°C. The supernatant was mixed with Protein A beads coupled to rabbit anti-GFP antibodies and rotated end-over-end at 4°C for 2 hours. Beads were rinsed twice in wash buffer (50 mM HEPES pH 7.4, 1 mM EGTA, 1 mM MgCl2, 300 mM KCl, 10% glycerol, 0.05% NP-40, 1 mM DTT, 10 μg/mL leupeptin/pepstatin/chymostatin). Beads were then washed 3 times in wash buffer by rotating for 5 minutes at 4°C. Bound protein was eluted with 100 mM glycine pH 2.6, precipitated by addition of 1/5th volume TCA at 4°C overnight. TCA precipitant was washed 3 times with -20°C acetone, dried using a speed vac, and stored at -20°C.

### Mass spectrometry sample preparation

Proteins were digested and cleaned up using a modified version of the S-trap protocol (Protifi, v4.7). TCA precipitant was resuspended in 23 µL 1x lysis buffer (5% SDS, 50 mM TEAB pH 8.5), denatured at 95°C for 10 minutes in the presence of 20 mM DTT. Samples were alkylated with 40 mM iodoacetamide for 30 minutes at room temperature then acidified to a final concentration of 2.5% v/v phosphoric acid. 6X volume of S-trap binding/wash buffer (165 µL) was added then loaded onto S-trap mini columns. Samples were spun at 4000x g for 30 seconds and washed 4 times with 150 µL S-trap binding/wash buffer by spinning at 4000x g for 30 seconds. After the last wash, the column was dried by spinning at 4000x g for 1 minute. Proteins on column were digested overnight at 37°C in a humidified incubator with 20 µl of 50 mM TEAB pH 8.5 containing 1 µg of trypsin. Peptides were eluted using 40 µl of 50 mM TEAB, 0.2% formic acid, then 50% acetonitrile. The eluted peptides were pooled, quantified using the Themo fluorescent peptide quantification kit, flash frozen in liquid nitrogen, then lyophilized.

### TMT labelling and peptide fractionation

1 µg trypsinized peptides were dissolved in 50 mM TEAB pH 8.5 and labelled using the TMT10plex Isobaric Labeling Reagent Set (Thermo Fisher Scientific, 90111). Each sample was labeled with TMT10plex reagents at a 20:1 label:peptide w/w ratio, for 1 hour at room temperature. TMT labeling reaction was quenched with 0.2% hydroxylamine for 15 minutes at room temperature. The samples were pooled on ice, flash frozen and lyophilized. Pooled TMT-labelled peptides were cleaned and fractionated using the Pierce High pH Reversed-Phase Peptide Fractionation Kit (Thermo Fisher Scientific, 84868) according to manufacturer’s instruction for TMT experiments. After fractionation, samples were flash frozen and lyophilized.

### Mass spectrometry data acquisition

Lyophilized peptides were resuspended in 0.1% formic acid to a final concentration of 250 ng/uL. Mass spectrometry was performed using Eclipse Orbitrap mass spectrometer equipped with a FAIMS Pro source connected to an Vanquish Neo nLC chromatography system. Our samples are separated using either a 25 cm analytical column (PepMap RSLC C18 3 µm, 100A, 75 µm). For all mass spectrometry experiments, peptides were separated at 300 nl/min on a gradient of 3–25% B for 57 min, 25–40% B for 17 min, 40– 95% B for 10 min, 95% B over 6min, and then an equilibration of the column to 3% B at the end of the run using 0.1% FA in water for A and 0.1% FA in 80% acetonitrile for B. The orbitrap and FAIMS were operated in positive ion mode with voltage of 2100 V; with an ion transfer tube temperature of 305°C, 4.2 l/min carrier gas flow, using standard FAIMS resolution and compensation voltages of –50 and –65 V. Full scan spectra were acquired in profile mode at a resolution of 120,000 (MS1) and 50,000 (MS2), with a scan range of 400–1400 m/z, custom maximum fill time (200ms), custom AGC target (300% MS1, 250% MS2), isolation windows of m/z 0.7, intensity threshold of 2x104, 2–6 charge state, dynamic exclusion of 60 seconds, and 38% HCD collision energy.

### Mass spectrometry analysis

Raw files were analyzed in Proteome Discoverer 2.4 (Thermo Fisher Scientific) to generate protein and peptide IDs using Sequest HT (Thermo Fisher Scientific) and the Homo sapiens protein database (UP000005640) with EGFP. The maximum missed cleavage sites for trypsin was limited to 2. Precursor and fragment mass tolerances were 10 ppm and 0.02 Da, respectively. The following post-translational modifications: dynamic oxidation (+15.995 Da; M), dynamic acetylation (+42.011 Da; N-terminus), dynamic Met-loss (–131.04 Da; M N-terminus), dynamic Met-loss+acetylation (–89.03 Da; M N-terminus), static carbamidomethyl (+57.021 Da; C), static TMT6plex (+229.163 Da; any N-terminus), and static TMT6plex (+229.163 Da; K). TMT 10plex isotope correction values were accounted for (Thermo Fisher; 90111 LOT# VK306786). Peptides identified in each sample were filtered by Percolator to achieve a maximum FDR of 0.01.

### Live cell fluorescence microscopy

Cells on glass bottom plates were incubated in 0.1 μg/mL hoescht for >30 minutes. For Aurora kinase inhibition experiments, cells on glass bottom plates were treated with 10 μM ZM447439 at 37°C for 2.5 hours. In the last 30 minutes before completing ZM447439 treatment, 0.1 μg/mL hoescht was added to the media. Images were taken on the Deltavision Ultra (Cytiva) system using a 60x/1.42NA objective. 8 μm images were taken with z-sections of 0.2 μm. All images presented were deconvolved and max projected. Images presented within the same panel were taken on the same day, camera settings, and scaled equivalently. Images with morphological markers such as DNA or tubulin were not all scaled equivalently.

### Image quantification

The relative mitotic GFP-DSN1 kinetochore intensities were quantified with a custom automated CellProfiler (v4.2.6, [36]) pipeline. The integrated intensity of a 5-pixel wide region surrounding each kinetochore was used to background subtract each measurement. Mitotic chromosomes were detected with the options: min/max diameter of 100 - 500, using the global minimum cross-entropy thresholding method. The size of the adaptive window is 50 without log transformation before thresholding. Puncta (or kinetochores) within chromosomes were segmented with the options: min/max diameter of 2 – 25, global robust background thresholding. The background was identified using Distance – N option and expanded 5 pixels from the identified kinetochore. The integrated EGFP intensity was measured within the kinetochore and background subtracted. Image quantification was performed on raw, non-deconvolved, maximally projected images.

### Competitive growth assays

Fluorescent mCherry HeLa cells were generated by infection with lentivirus carrying mCherry. Cells at ∼90% confluency were spinfected with lentivirus in the presence of 10 μg/mL polybrene. After 16 hours, the media was swapped. 24 hours post transduction, cells were moved into a new plate. After another 24 hours in culture, mCherry positive HeLa cells were bulk sorted. Equal amounts of control HeLa mCherry cells were mixed with uncolored DSN1^Δexon3^ cells. 3 independent DSN1^Δexon3^ clones were tested in the competitive growth assay. The population of mCherry:uncolored cells were monitored every few days using the BD LSRFortessa Cell Analyzer (BD Biosciences). Flow cytometry data was analyzed using FlowJo (v10.7.1).

### DSN1 Δintron2 mouse ESC and model generation

px330-sgRNA plasmids along with, DSN1 **Δ**intron 2 recombination plasmid, and mCherry-expressing plasmids were co-transfected into pre-plated v6.5 mouse ESC cells (without any feeder MEFs) using Lipofectamine 2000, according to the manufacturer’s protocol. mCherry positive ESCs were bulk sorted 48 hours post-transfection and sparsely cultured on MEF feeder gelatin-coated plates. Single clones were isolated and genotyped by PCR. To genotype, cells were lysed in DirectPCR lysis reagent (Viagen, 301-C) with proteinase K (0.2 µg/mL) overnight at 55°C then proteinase K was inactivated at 85°C for 90 minutes. Clones were screened using oligos (5’-CTGTGATGCCTGGGTGGAAGGG-3’ and 5’-AGGAGCTAAGATAGGTTCTGTAGACCG-3’) flanking intron 2 such that removal of intron 2 would result in a shorter PCR product. The PCR product was sequenced to ensure accurate homology-directed repair. Blastocyst injections were performed using (C57BL/6xDBA) B6D2F1. 6–8-week old B6D2F1 females were hormone primed by an intraperitoneal injection of pregnant mare serum gonadotropin (PMS, EMD Millipore) followed 46 hours later by an injection of human chorionic gonadotropin (hCG, VWR). Embryos were harvested at the morula stage and cultured in a CO2 incubator overnight. On the day of the injection, groups of blastocysts were placed in drops of M2 medium. Approximately ten mESC DSN1 Δintron2 cells were injected into the blastocoel cavity of each embryo. Approximately 20 injected blastocysts were transferred to each recipient female. Chimera mice were mated to C57BL/6 the resulting pups were PCR screened for the DSN1 Δintron2 mutation. Animals containing the DSN1 Δintron2 edit was then back crossed at least 10 times prior to any experimentation. Animal experiments performed in this study were approved by the Massachusetts Institute of Technology Committee on Animal Care (0820-020-23).

### Mouse husbandry

For mouse generation and fertility assays, mice were housed in a 12-12 h light/dark cycle with constant temperature and food at the Whitehead Institute. The Massachusetts Institute of Technology Department of Comparative Medicine provided daily cage maintenance and health checks. Mouse generation and care were approved by the Massachusetts Institute of Technology Committee on Animal Care (0820-020-23).

For oocyte and embryo DSN1 experiments, all animals were maintained in accordance with the guidelines and policies from the Institutional Animal Use and Care Committee at Rutgers University (Protocol# 201702497). Mice were housed in a room programmed for a 12-hour dark/light cycle and constant temperature (between 70-74°F), humidity (50%) and with food and water provided ad libitum.

### Metaphase II egg and embryo collections and embryo development

For egg collection, females 6-16 weeks old were hormonally stimulated with 5 I.U. of pregnant mare serum gonadotropin (PMSG) (BioVendor R&D; RP1782725000) to promote follicle growth. 48h post PMSG injection, females were injected with 5 I.U of human chorionic gonadotropin (hCG) (Sigma-Aldrich #CG5) to induce ovulation, and 12 h hours later, ovulated metaphase II eggs were collected from ampullas. The ampullas were dissected in MEM supplemented with 3mg/ml Hyaluronidase (Sigma; H3506).

To collect embryos, WT and mutant females (6-16 weeks) were hormonally stimulated as above. After hCG females were mated with WT or mutant males. There were four mating combinations: C57BL/6 female X C57BL/6 male; DSN1 Δintron2/Δintron2 female X C57BL/6 male; and DSN1 Δintron2/Δintron2 female X DSN1 Δintron2/Δintron2 male. 20 hours after hCG, 1 cell embryos were collected from ampullas and culture in KSOM embryo culture media (Millipore; MR.106-D) in an atmosphere of 5% CO2 in air at 37°C. Images of embryos were collected 48h and 96 h post-hCG under a bright field microscope EVOS FL Auto Imaging System (Life Technologies) using a 20x objective.

### Immunofluorescence

HeLa cells were seeded on poly-L-lysine coated coverslips and fixed in PHEM + 4% formaldehyde + 0.25% Triton X-100 at room temperature for 15 minutes. For Aurora kinase inhibition experiments, cells on coverslips were treated with 10 μM ZM447439 at 37°C for 2.5 hours prior to fixation. Following fixation, cells were washed 3 times with PBS + 0.1% Triton X-100 and blocked in AbDil for 30 minutes - 1 hour. Following blocking, cells were stained at room temperature for 1 hour with α-DSN1 (1 µg/mL, [10]), α-NDC80 bonsai (1 µg/mL, [37]), α-alpha tubulin (1:1000, Sigma-Aldrich, T9026), α-centromere antibody (1:300, Antibodies Inc, 15-234-0001), α-phospho-histone H2A.X (Ser139) antibody (1:1000, Millipore, JBW301) diluted in AbDil. Cells were washed with PBS + 0.1% Triton X-100, 3 times then incubated in secondary antibody diluted in AbDil for 1 hour at room temperature. After secondary, cells were stained in hoescht for 15 minutes at room temp, then washed 3 times with PBS + 0.1% Triton X-100 before mounting in PPDM (0.5% p-phenylenediamine and 20 mM Tris-Cl, pH 8.8, in 90% glycerol) and sealed with nail polish. Images were taken on the Deltavision Ultra (Cytiva) system using a 60x/1.42NA objective. 8 μm images were taken with z-sections of 0.2 μm. All images presented were deconvolved and max projected. Images within the same panel were taken on the same day, with same camera settings, and scale equivalently. Images with morphological markers such as DNA or tubulin were not all scaled equivalently.

The ovulated eggs were immunoassayed as described [38]. Briefly, eggs were fixed in phosphate-buffered saline (PBS) containing 2% paraformaldehyde (PFA) (Sigma; P6148) at room temperature for 20 minutes. Then, oocytes were incubated for 20 minutes in a permeabilization solution at room temperature (PBS containing 0.1% (vol/vol) Triton X-100 and 0.3% (wt/vol) BSA). Eggs were blocked for 10 minutes (0.3% BSA containing 0.01% Tween in PBS). Immunostaining was performed by incubating cells in primary antibody, Pericentrin (PCNT) (mouse, 1:100; BD Biosciences, #611814) and Alpha Tubulin-conjugated 488 (Rabbit, Cell Signaling; S063S) for 1 hour in a dark, humidified chamber at room temperature followed by 3 consecutive 10-minute incubations in blocking buffer. After washing, the secondary antibody, Alexa anti mouse-568 (Invitrogen; A10037) was diluted 1:200 in blocking solution and incubated with the cells for 1 h at room temperature. After washing, the cells were mounted in 10 μL VectaShield (Vector Laboratories, #H-1000) containing 4′, 6-Diamidino-2-Phenylindole, Dihydrochloride (DAPI; Life Technologies #D1306; 1:170).

Egg images were captured using a Leica SP8 confocal microscope equipped with a 40x, 1.30 NA oil immersion objective. Optical Z-stacks of 1.0 μm step with an optical zoom of 4.0.

### Fertility trials

Sexually mature (6-8 weeks old) females were continuously mated to male mice (6-8 weeks old). Mating’s were continued for 6 months and cages were checked regularly for pups.

## Supplemental Figures

**Figure S1.**
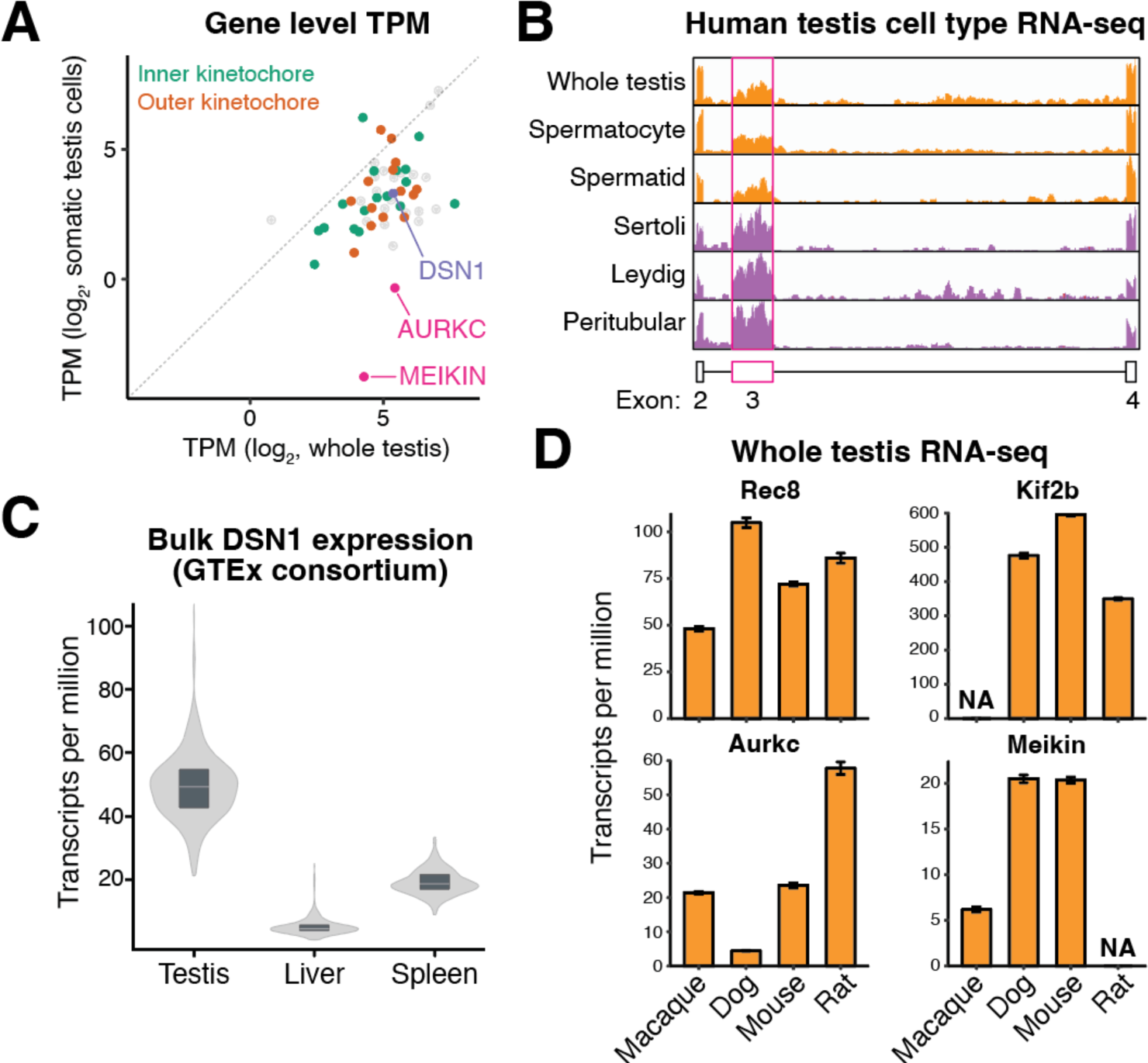
Expression analysis of DSN1 and conserved localization of mouse DSN1. (**A**) Comparison of gene level mRNA sequencing from mitotic HeLa and whole human testis for kinetochore proteins. (**B**) Genome browser track of mRNA sequencing data showing DSN1 exon expression in various human testis cells. Exon 3 reads are highlighted in the magenta box. The relative reads mapped to exon 3 are depleted in any tissue with meiotic cells. (**C**) Human DSN1 mRNA expression from the GTEx consortium in testis, liver, and spleen. Total DSN1 levels are decreased in liver cells, consistent with RT-PCR analysis in mouse tissues (Fig. 1H). (**D**) Transcripts per million for Rec8, Kif2b, Aurkc, and Meikin in whole testis from indicated species. NA indicates that this gene was not detected in the downloaded genome annotation. Error bars indicate standard error of the mean and N = 8 biological replicates for Macaque, Dog, and Mouse. N = 7 biological replicates for Rat testis.

**Figure S2.**
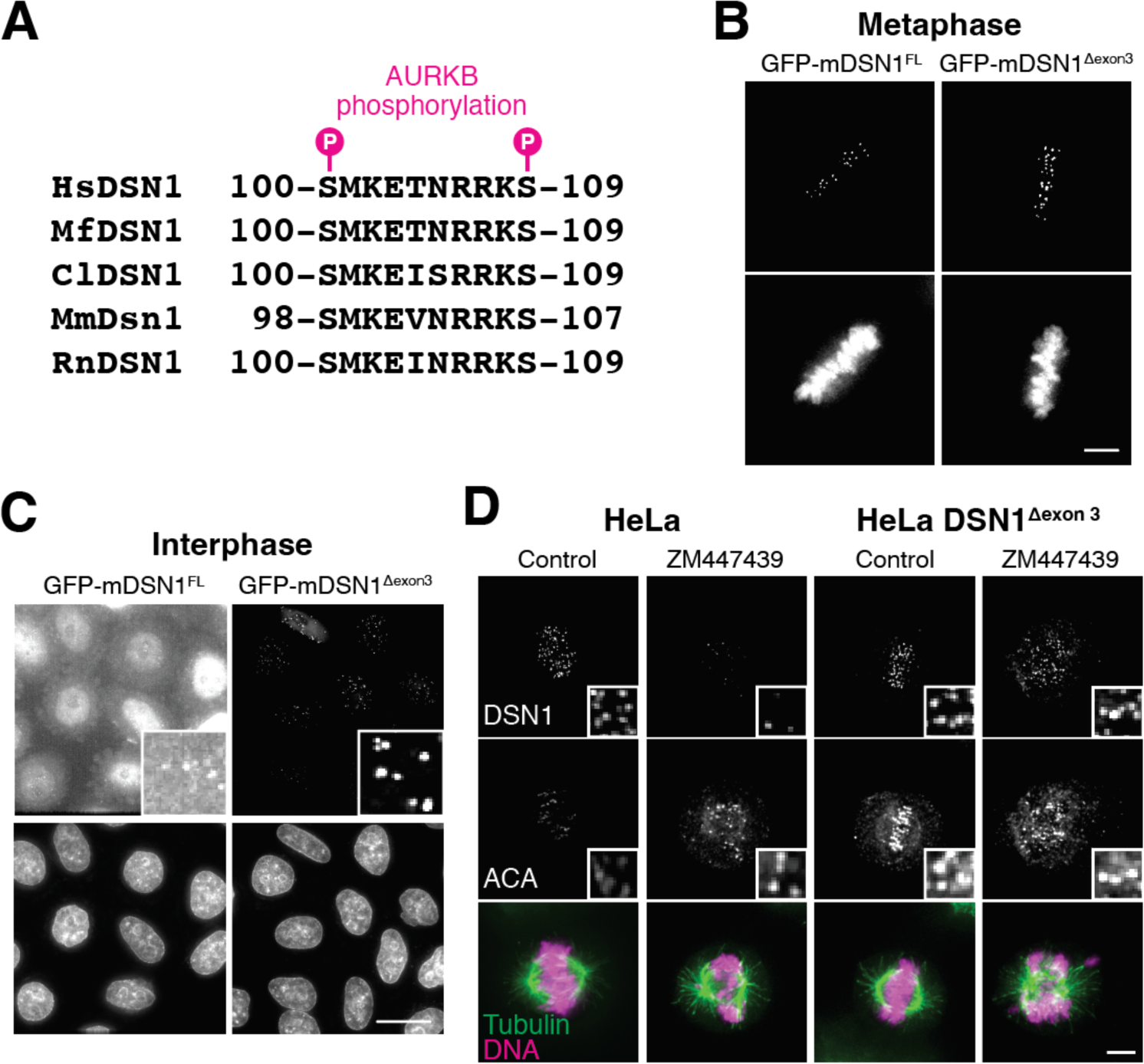
Regulatory control of human DSN1 is conserved in mouse DSN1. (**A**) Conservation of DSN1 phosphorylation sites across humans, macaque, dog, mice, and rats. (**B**) Live cell imaging showing the localization of ectopically expressed GFP-mouse DSN1 full length and mouse GFP-DSN1^Δexon3^. Similar to the human constructs, both constructs are able to localize to centromeres during interphase. Scale bar represents 5 µM. (**C**) Live cell imaging of GFP-mouse DSN1 full length and GFP-mouse DSN1^Δexon3^ during interphase. GFP-mouse DSN1^Δexon3^ is still able to localize to centromeres. Scale bar represents 20 µM. (**D**) Immunofluorescence images of DSN1 localization in control and DSN1^Δexon3^ cell lines in the presence or absence of the Aurora kinase inhibitor ZM447439. Scale bar represents 5 µM.

**Figure S3.**
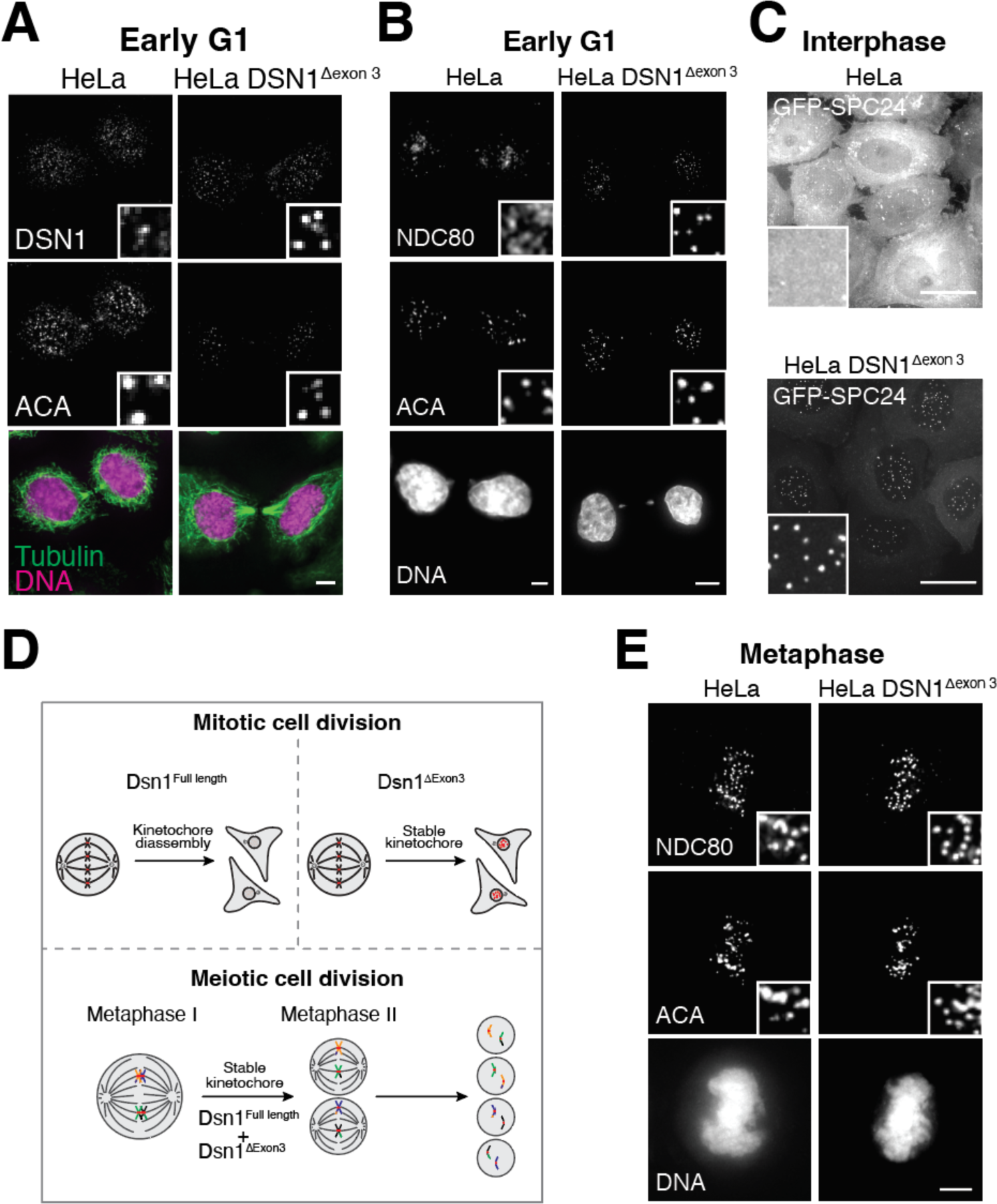
Analysis of HeLa cells lacking endogenous DSN1 exon 3. (**A**) Immunofluorescence images of DSN1 in control and DSN1^Δexon3^ cell lines during early G1. Scale bar represents 5 µM. (**B**) Immunofluorescence images of NDC80 in control and DSN1^Δexon3^ cell lines during early G1. Scale bar represents 5 µM. (**C**) Live cell imaging of GFP-SPC24 during interphase in control and DSN1^Δexon3^ cell. GFP-SPC24 is able to localize to interphase centromeres in cells lacking DSN1 exon 3. Scale bar represents 20 µM. (**D**) Immunofluorescence images of NDC80 in control and DSN1^Δexon3^ cell lines during metaphase. Scale bar represents 5 µM. (**E**) Schematic model of the role of DSN1 splicing in mitotic cells and proposed role of DSN1 splicing during meiosis. Full length DSN1 is removed from centromeres after chromosome segregation and removal of DSN1 exon 3 results in persistent centromere localization. During meiosis, DSN1^Δexon3^ is expressed, which may allow for constitutive centromere localization.

**Figure S4.**
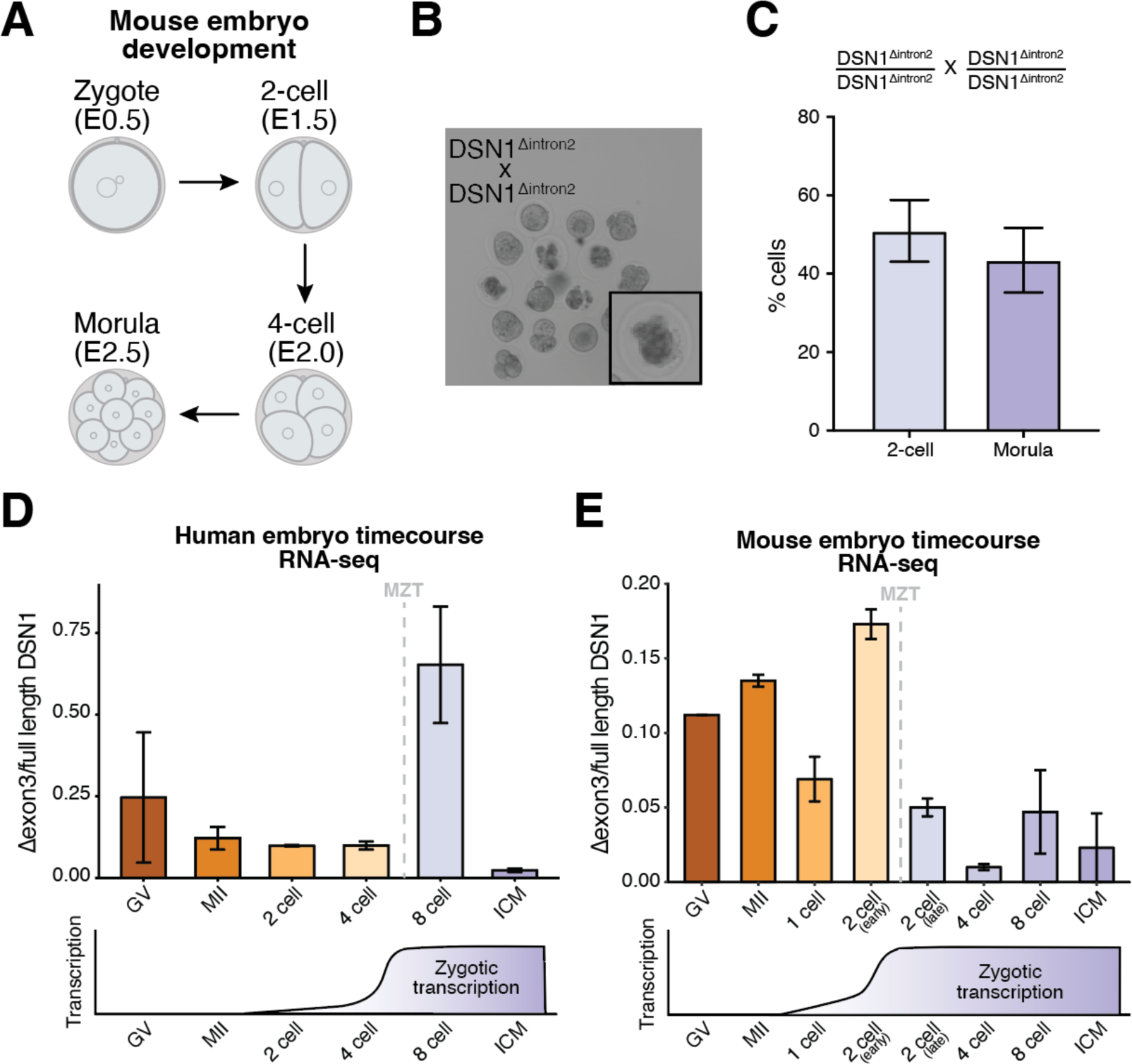
Development of DSN1 mouse embryos and evidence for early zygotic expression of DSN1 exon3. (**A**) Schematic outline of mouse embryo development. (**B**) Representative images of 8 cell embryos (morula) from a DSN1Δintron2/Δintron2 x DSN1Δintron2/Δintron2 mating. (**C**) Quantification of the number of 2 cell and morula embryos from a DSN1Δintron2/Δintron2 x DSN1Δintron2/Δintron2 mating. N = 3. (**D**) DSN1^Δexon3^/DSN1^full^ ^length^ mRNA expression across human embryo development. DSN1^Δexon3^ is transiently expressed near the onset of zygotic transcription. Error bars indicate standard error of the mean and N = 3. (**E**) DSN1^Δexon3^/DSN1^full^ ^length^ mRNA expression across mouse embryo development. DSN1^Δexon3^ is transiently expressed near the onset of zygotic transcription. Error bars indicate standard error of the mean and N = 3.

